# Oropouche virus causes acute hepatitis in mice controlled by Type I interferons

**DOI:** 10.64898/2026.04.16.718884

**Authors:** Cade E. Sterling, Rachael E. Rush, Jackson J. McGaughey, Brooke A. Snow, Alexandra J. Benton, W. Paul Duprex, Gaya K. Amarasinghe, Satdarshan P. Monga, Amy L. Hartman

**Author notes:** Address correspondence to Amy L. Hartman.

## Abstract

Oropouche virus (OROV), a member of the *Peribunyaviridae* family endemic to South America, is a current public health threat. The recent OROV outbreak driven by a novel reassortant has caused a dramatic increase in cases in 2024 (13,785 in Brazil, versus only 261 from 2015-2022) with sustained levels of transmission in 2025. Previously underreported outcomes have been recognized including miscarriage, microcephaly, encephalitis, and death. OROV lethality in humans has been attributed to severe coagulopathy with liver involvement, and epidemiological data suggests acute hepatitis occurs in mild cases of Oropouche fever, highlighting the underrecognized role of the liver in OROV pathogenesis. We present two discrete mouse models of OROV hepatic disease — a lethal model that recapitulates the severe coagulopathy seen in fatal human cases and a model of self-resolving acute hepatitis which recapitulates mild human disease. OROV causes focal hepatic necrosis in mice, which progresses to massive necrosis and death when the Type I interferon receptor is antagonized. Additionally, we found a contemporary OROV isolate is less pathogenic in mice than a historic prototypical strain. These studies enhance our understanding of OROV pathogenesis and pave the way for potential therapeutic development and evaluation.

**Importance:** The disease burden of Oropouche fever has been underrecognized and underreported, as highlighted by the increased testing seen in the ongoing outbreak. Specifically, the role of the liver in Oropouche virus pathogenesis has been neglected. Given the ongoing outbreak and increase in severe disease manifestations, there is a present need to understand Oropouche virus pathogenesis and test potential therapeutics. The mouse models of Oropouche-induced hepatitis presented here provide a means to understand how Oropouche virus causes liver damage in both a lethal and sublethal context. These models will be useful for the preclinical evaluation of vaccines and therapeutic treatments. Additionally, we compare the pathogenicity of a historical Oropouche virus isolate to a contemporary human isolate in a lethal mouse model. This represents an additional step towards understanding whether the circulating Oropouche virus isolates are uniquely more pathogenic or if increased testing has highlighted previously unreported outcomes.

## Introduction

Oropouche virus (OROV; class *Bunyaviricetes*, family *Peribunyaviridae*, Simbu serogroup) causes sporadic human outbreaks, primarily in the Amazon Basin, where spillover occurs from sloths, non-human primates, and birds via *Culicoides* midges or, to a lesser extent, *Culex* mosquitos.^1^ In humans, Oropouche fever most often presents as a self-limiting febrile illness characterized by headaches, myalgia, joint pain, and gastrointestinal distress, though occasional cases of sever neurologic disease have been reported.^2,3^

The largest recorded OROV outbreak is currently ongoing with over 16,000 confirmed cases in the Americas in 2024 alone and a resurgence of cases in 2025 at similar levels.^4–7^ Genomic analyses of OROV cases from August 2022 to February 2024 discovered that this outbreak coincided with the emergence of a novel reassortant lineage of OROV.^8^ Brazil endured the largest disease burden with 831 reported cases in 2023 and 13,785 cases in 2024 (compared to only 216 total cases between 2015-2022).^6,9,10^ Imported travel-associated OROV cases were found in non-endemic regions including the United States, Italy, Spain, and Germany.^11–13^ Moreover, there has been OROV emergence and ongoing transmission in Cuba, highlighting the ability of OROV to establish endemic transmission in new regions.^14,15^ Notably, competent vectors are found in the United States, Canada, and Europe.^12,16–19^

Beyond increased transmission and reported cases, the 2023-2025 outbreak resulted in recognition of previously unreported forms of clinical disease including adverse pregnancy outcomes (i.e., vertical transmission, fetal loss, microencephaly), serve neurologic outcomes in adults (i.e. Guillain-Barré syndrome), hemorrhagic fever, and death.^4,11,14,20–24^ Prior to this outbreak, there were no known cases of lethal Oropouche fever. It is unclear whether the reassortant strain currently circulating is uniquely more virulent or if the increase in cases and testing has revealed previously undocumented severe outcomes. Taken together, the threat OROV poses to the Americas is apparent, reflected by the fact that the Pan American Health Organization (PAHO; a World Health Organization regional office) issued a Public Health Risk Assessment rating the regional risk as high.^5^ Consequently, there is a present need to understand the severe outcomes of Oropouche fever and test potential therapeutics. To this end, the development of relevant animal models is paramount.

The cause of death in the first two lethal human cases of Oropouche fever, which occurred in otherwise healthy individuals, was attributed to severe coagulopathy with liver involvement.^4^ The third lethal case resulted from severe coagulopathy, respiratory distress, and renal failure; liver histology showed signs of necrotic foci, steatosis, and severe hepatic damage.^25^ More recent case studies have also documented Oropouche-associated hepatitis in otherwise healthy patients with elevated hepatic transaminases.^26,27^ While it has been previously reported that mild cases of Oropouche fever can coincide with mild liver injury,^3^ the role of the liver in OROV pathogenesis has been largely unexplored.

Here, we report two mouse models of OROV hepatic disease — a sublethal and lethal model. In the sublethal model, immunocompetent mice develop a transient, acute self-resolving hepatitis characterized by elevated liver enzymes, necrotic foci, and robust OROV replication in the liver. This likely represents a model of mild human cases of Oropouche fever. In the lethal model, OROV susceptibility is induced by administration of an anti-Ifnar1 antibody which results in mice experiencing lethal hepatic disease characterized by severe coagulopathy, sinusoidal congestion, and hemorrhagic necrosis, much like the lethal human cases of Oropouche fever. The development and characterization of these models will be useful for testing vaccines, therapeutics, and other medical countermeasures against OROV.

## Results

### OROV induces non-lethal transient acute hepatitis in immunocompetent mice

Immunocompetent adult laboratory mice do not generally show visible signs of disease following OROV infection from peripheral inoculation routes (intradermal, subcutaneous, or intraperitoneal). Indeed, infection of immunocompetent C57BL/6 mice with recombinant OROV BeAn19991 by footpad infection (mimicking a mosquito bite) resulted in sublethal infection with no clinical signs of disease including a lack of weight loss (Supplementary Figure S1). This contrasts with intranasal inoculation of mice which succumb to infection (Supplementary Figure S1). Despite lack of outward signs of disease or lethality following footpad injection, we sought to determine whether OROV replicates systemically in immunocompetent mice. We infected C57BL/6 mice with 10^6^ PFU of the prototypical OROV strain BeAn19991 via the footpad and euthanized a subset of animals at 1 (n=11), 3 (n=15), and 5 (n=11) days post infection (dpi; Figure 1A). Upon necropsy, tissues were harvested for measurement of viral load and histology. At 3 dpi, OROV-infected mice had transiently elevated levels of alanine transaminase (ALT), but not alkaline phosphatase (ALP) or other parameters (Figure 1B, C; Supplementary Figure S2); these blood chemistry results are indicative of mild hepatocellular damage. Median blood chemistry parameters were within the normal range (95% CI for C57BL/6J mice) at 1 and 5 dpi.^28^ While the elevation at 3 dpi is not statistically significant, the median elevation seen is clinically relevant (7-fold increase over uninfected controls and over double the upper limit of normal). Infectious OROV was detectable in the liver as early as 1 dpi, with peak titers at 3 dpi (10^3^–10^8^ PFU/g); by 5 dpi, most mice returned to undetectable levels of infectious virus (Figure 1D). We found no infectious virus in the spleen, kidney, lung, or brain at any time point, and low to moderate levels of viral RNA were detected in these tissues (Supplementary Figure S3). Taken together, these data suggest that inapparent OROV infection is hepatotropic in C57BL/6 mice and causes a self-resolving acute hepatitis.

**Figure 1.**
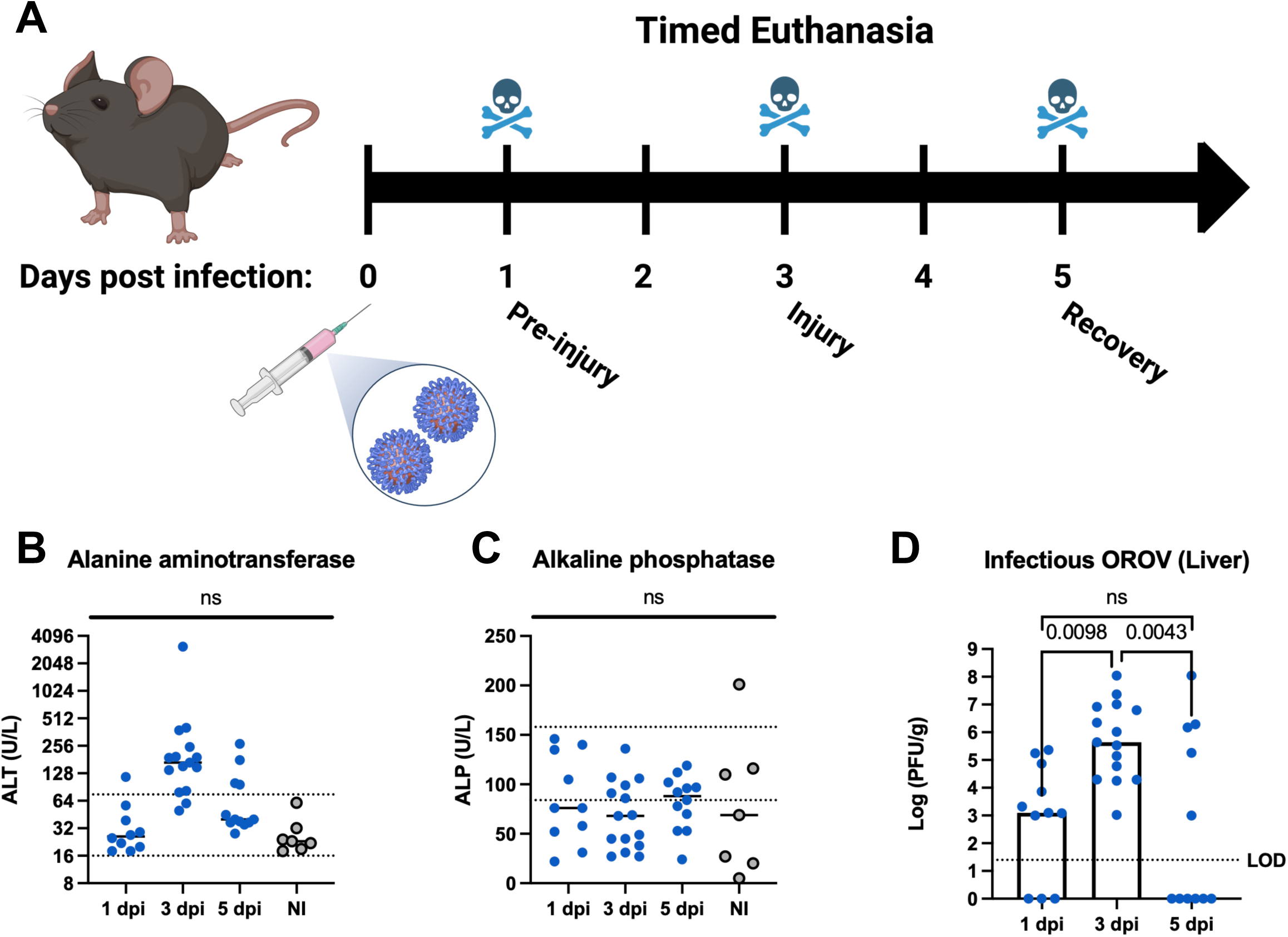
OROV induces transient acute liver injury in immunocompetent mice. (A) Schematic of OROV acute hepatis model. Mice were infected with 10^6^ PFU OROV via footpad injection and a subset were euthanized at 1 (n=11), 3 (n=15), and 5 (n=11) dpi. (B) Blood alanine aminotransferase (ALT) and (C) alkaline phosphatase (ALP) levels at indicated timepoints post infection or uninfected controls (n=7). (D) Infectious OROV liver titers by plaque assay at indicated timepoints post infection. Each data point represents one mouse across three experiments; bars represent median value. Dotted lines indicate either normal (95 confidence interval) range or limit of detection (LOD). Statistical significance was determined by ordinary one-way ANOVA and Tukey’s multiple comparisons test. P value shown or ns (not significant) if p > 0.05. Uninfected controls labeled “NI” (not infected).

Given that we found considerable viral titers in the liver along with elevated liver enzymes at 3 dpi, we examined liver tissue for histological damage. Sections of the left lateral lobe were stained by H&E which reveled discrete inflammatory foci (black arrows) throughout the lobule (Figure 2A). These foci were characterized by a breakdown in cellular architecture, the accumulation of eosin staining (either blood or cytoplasmic accumulation), and a relatively high number of immune infiltrates (the small nuclei seen in the foci; Figure 2A). The foci were often surrounded by abnormally swollen hepatocytes (blue arrows) which is a nonspecific marker of injury (Figure 2A). To determine if these foci were also areas of cell death, we performed TUNEL staining and found that inflammatory foci were highly TUNEL positive (Figure 2B). The degree of ALT elevation at 3 dpi varied between animals (Figure 1B), and the total area of TUNEL staining across the lobule generally reflected the relative ALT levels (i.e., animals with higher blood ALT levels had more extensive hepatic necrosis; Supplementary Figure S4). By immunofluorescent staining, these foci were also positive for OROV N antigen (Figure 3C), suggesting they are sites of viral infection. It is unclear whether the accumulation of OROV antigen is the result of direct infection of hepatocytes that formerly occupied that space or if the infiltrating cells are infected themselves. Based on histopathologic examination, necroinflammatory foci appear to be stochastically distributed across the liver (Figure 2); however, there may be some preference for midzone (zone 2) and pericentral zone (zone 3) infection as judged by the relationship of these foci to portal triads or central veins. Further investigation into OROV zonal preference is warranted, as the observations here are similar to a human case which reported focal infection with a preference for zone 2 and 3 in the patient’s liver.^25^

**Figure 2.**
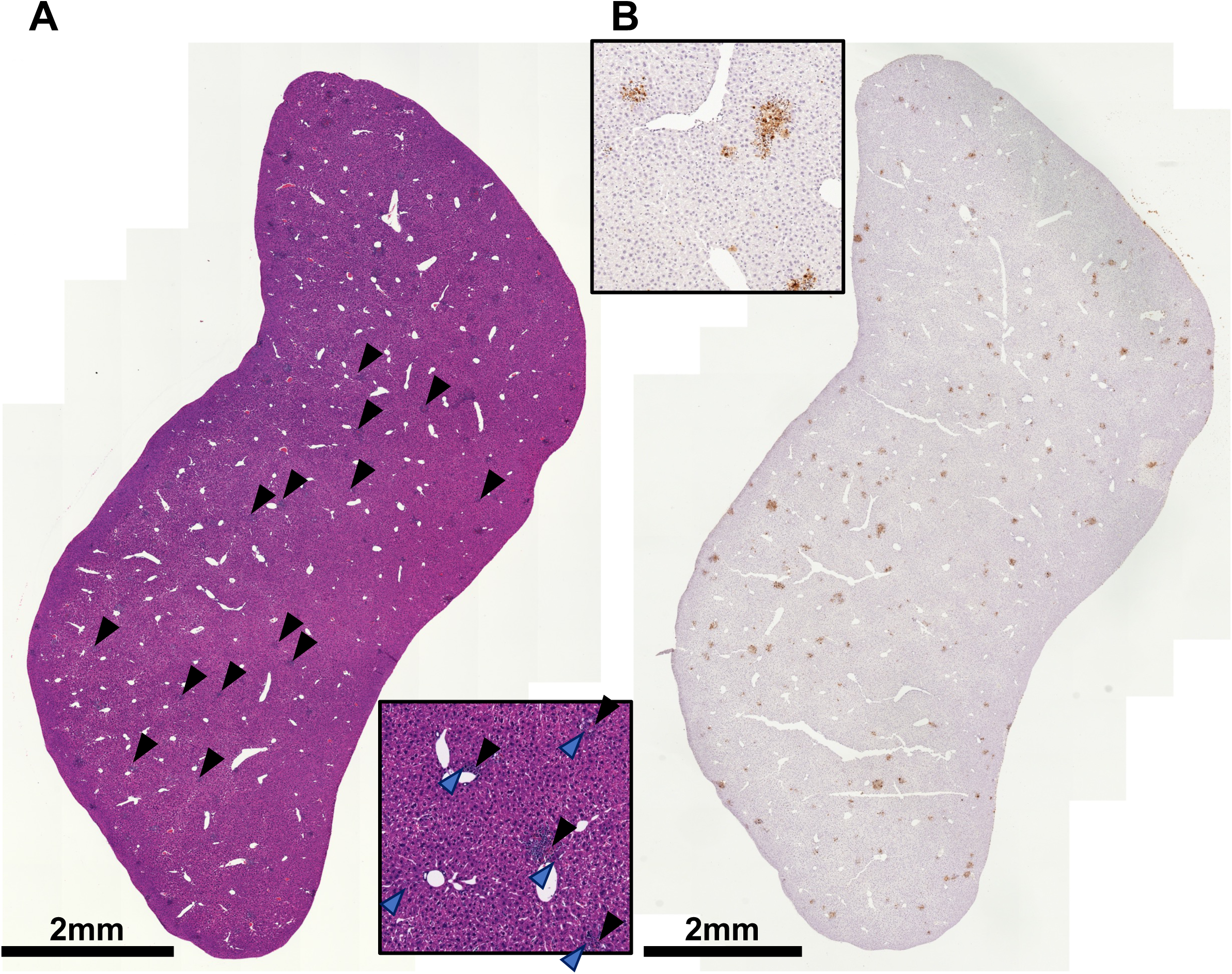
Histopathology reveals OROV-induced necroinflammatory foci across the entire lobule. (A) H&E and (B) TUNEL staining of a representative FFPE liver left lateral lobe from a mouse euthanized 3 days post infection (10^6^ PFU inoculation dose; total n=15, 3 independent experiments). Composite images and a representative 20X ROI (594×594 µm) are shown. Black arrows mark necroinflammatory foci; blue arrows mark regions of swollen hepatocytes. Not all features are marked.

**Figure 3.**
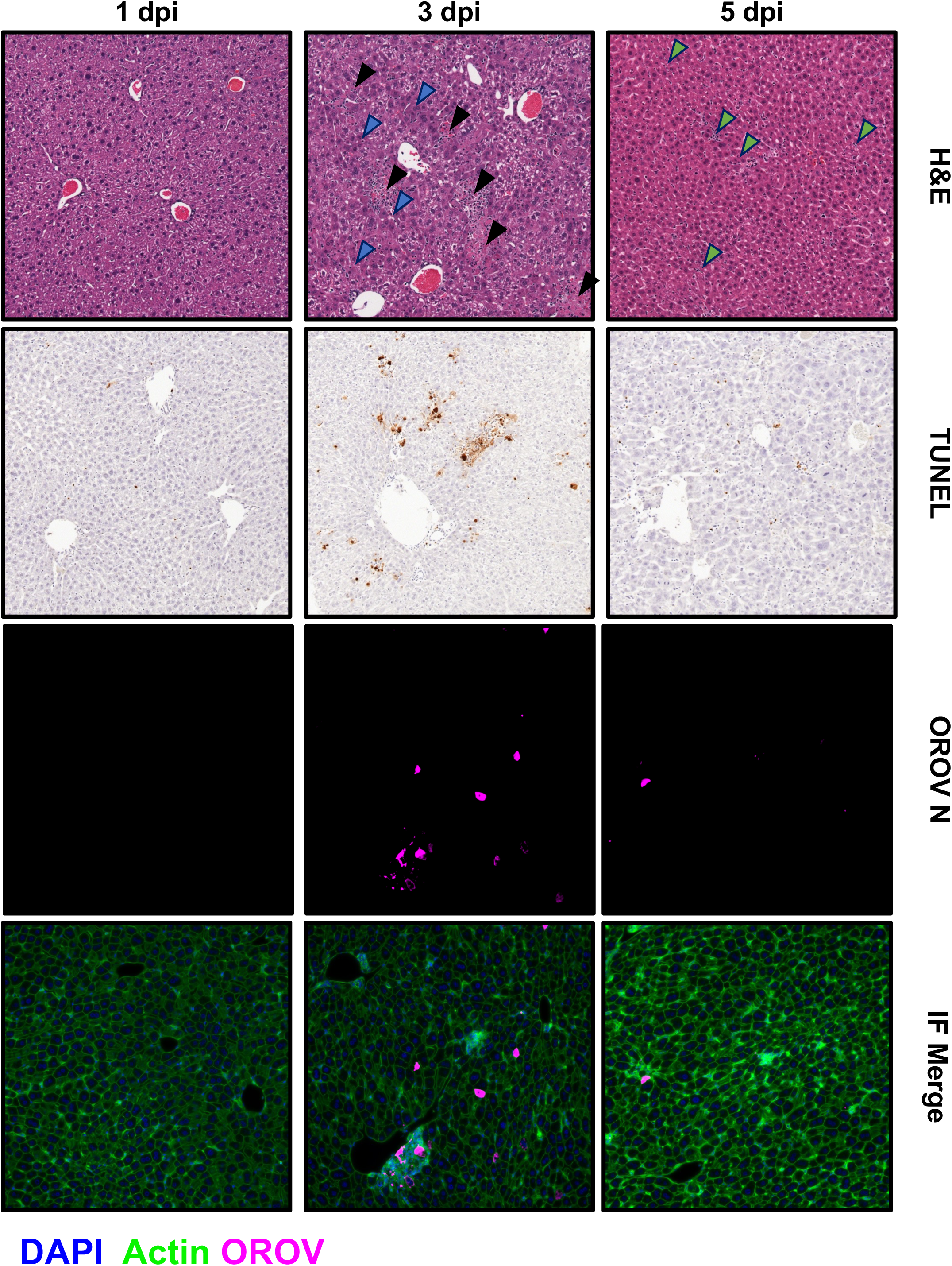
OROV-induced hepatic injury is transient. Representative 20X ROIs of H&E (594×594 µm), TUNEL (594×594 µm), and OROV N (670×670 µm) staining at 1 (n=11), 3 (n=15), and 5 (n=11) dpi following infection with 10^6^ PFU OROV via footpad injection. H&E and TUNEL staining from FFPE left lateral lobes; OROV N staining from cryo-embedded and sectioned right medial lobes. Labels indicate timepoint and stain. In H&E images, black arrows mark necroinflammatory foci; blue arrows mark regions of swollen hepatocytes; green arrows mark non-focal immune infiltrates. Not all features are marked.

We also examined whole liver lobules for histological damage at 1 and 5 dpi. Generally, the liver at 1 dpi was indicative of pre-injury whereas recovery was ongoing at 5 dpi (Figure 3; Supplementary Figure S5). While low levels of infectious virus were detected in the liver at 1 dpi (Figure 1C), there is a lack of histopathological damage and OROV antigen at this time (Figure 3). By 5 dpi, there was less infectious virus (Figure 1C), OROV antigen, and the necroinflammatory foci had resolved; however, there were still observable histopathological changes (Figure 3). Specifically, the recovering livers (5 dpi) showed an overall increase in infiltrating immune cells dispersed throughout the liver (green arrows), suggestive of ongoing repair. Notably, there was a lack of focal damage or accumulation of immune cells in foci at 5 dpi (Figure 3). While there was some increased TUNEL staining at 5 dpi compared to 1 dpi and uninfected controls, it is remarkably less than 3 dpi, further suggesting ongoing repair (Figure 3; Supplementary Figure S5, S6, S7).

### Anti-IFNAR1 antibody treatment induces OROV susceptibility in wildtype mice

Using an anti-mouse IFNAR1 antibody to transiently immunosuppress mice and induce susceptibility to disease is a well-established practice for other viral pathogens.^29–32^ The MAR1-5A3 clone mAb (hereafter referred to as MAR1 Ab) binds the mouse IFNAR-1 protein and prevents dimerization with IFNAR-2, thereby preventing recognition of Type I interferons by the Type I IFN receptor complex (INFAR-1/2 heterodimer).^33,34^ Here, we administered 500 µg of MAR1 Ab (or isotype control) 1 day before infection (−1 dpi) with OROV BeAn19991 via footpad infection. Mice were monitored for survival until 28 dpi (Figure 4A). Following MAR1 treatment, we found near uniform lethality (95%) by 5 dpi at a dose of 10^6^ PFU OROV (Figure 4B). At lower viral infection doses (10^5^, 10^4^, 10^3^ PFU), mice succumb to disease between 3 and 7 dpi in a dose-dependent manner with few survivors (Figure 4B). We saw no lethality in isotype control treated animals infected with 10^6^ PFU OROV, which was expected based on the existing literature and our previous findings (Supplementary Figure 1). MAR1-treated mice presented normally prior to rapid onset of disease, which was characterized by listlessness, hunching, piloerection, and abnormal breathing. Symptomatic mice quickly progressed to being unresponsive. Mice euthanized prior to succumbing to disease had high infectious OROV titers in gross liver homogenate (10^5^-10^8^ PFU/g; Figure 4C), which correlated with high levels of viral RNA (Figure 4D). In MAR1-treated mice, infectious virus and viral RNA were also detected in the spleen, kidney, lung, and brain at endpoint (i.e., time of euthanasia due to severe disease; Figure 4E, 4F). Infectious titers in the liver were higher than any other tested tissues.

**Figure 4.**
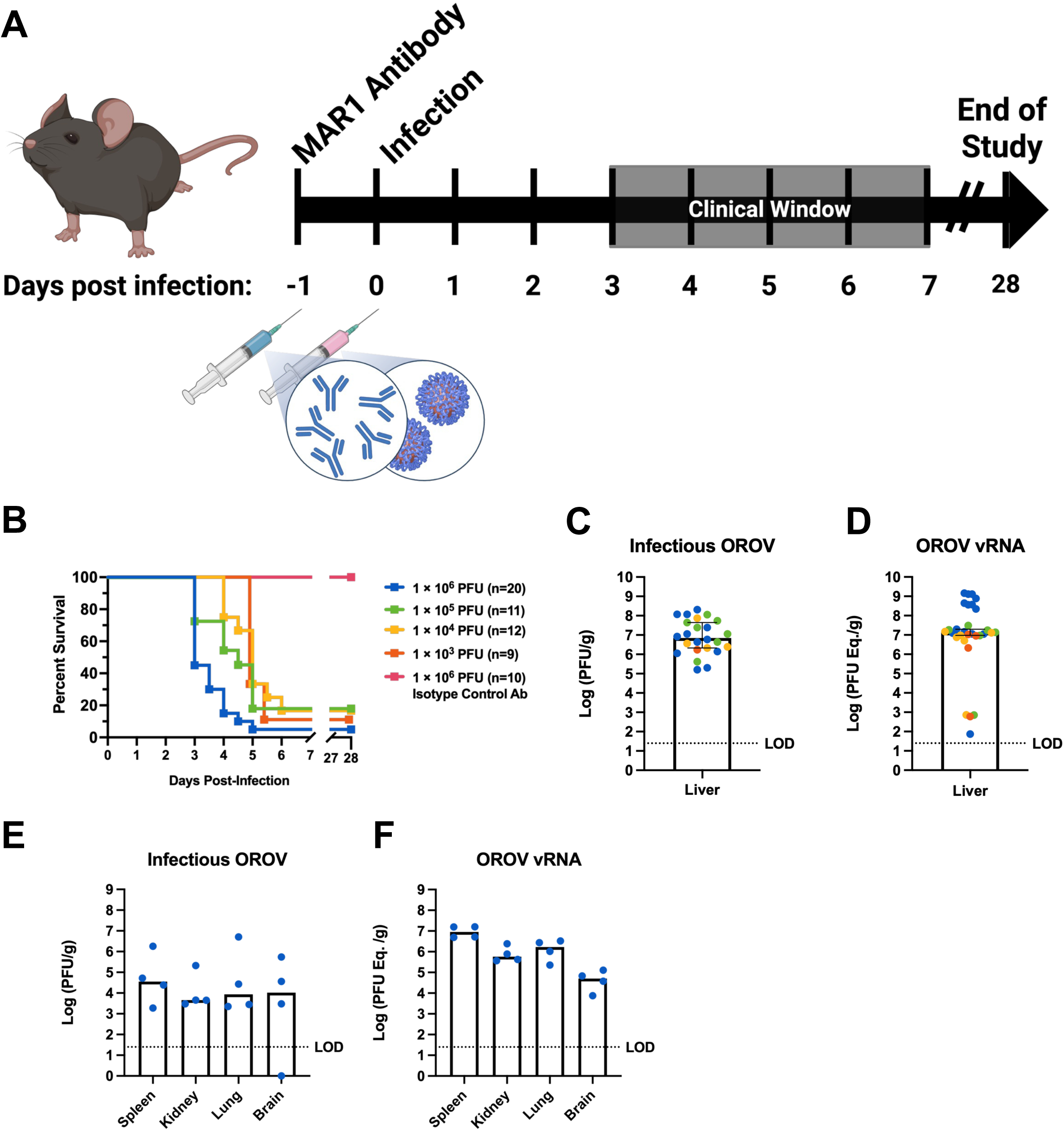
MAR1 antibody treatment induces OROV susceptibility in mice. (A) Schematic of lethal OROV MAR1 model. Mice were treated with 500 µg MAR1 Ab 1 day prior to OROV infection via intradermal footpad injection. (B) Combined survival curve from 5 independent experiments. (C) Infectious OROV titers in liver homogenate from tissues collected at endpoint. Figure excludes samples unquantifiable due to degradation. Symbols are color coded based on infection dose. (D) Viral RNA in liver homogenate from tissues collected at endpoint quantified by RT-qPCR. (E) OROV titers by plaque assay or (F) RT-qPCR in spleen, kidney, lung, and brain tissues from a subset of mice (n=4) at endpoint. Bars represent median values. Error bars represent 95% confidence interval. Dotted lines indicate limit of detection (LOD).

Histopathologic examination of mouse livers at endpoint revealed fulminant hepatitis with massive hepatic necrosis across the whole lobule even at the lowest tested infection dose (10^3^ PFU; Figure 5A). The cellular architectural damage, severe coagulopathy, sinusoidal congestion, hemorrhagic necrosis, and large increase in infiltrating immune cells was consistent across all animals that succumbed to OROV disease (Figure 5B). This pattern of pathology was observed in both mice euthanized at a humane endpoint and found dead on arrival (Figure 5C). For mice that recover from infection, there were no signs of histopathologic damage at 28 dpi (Supplementary Figure S8).

**Figure 5.**
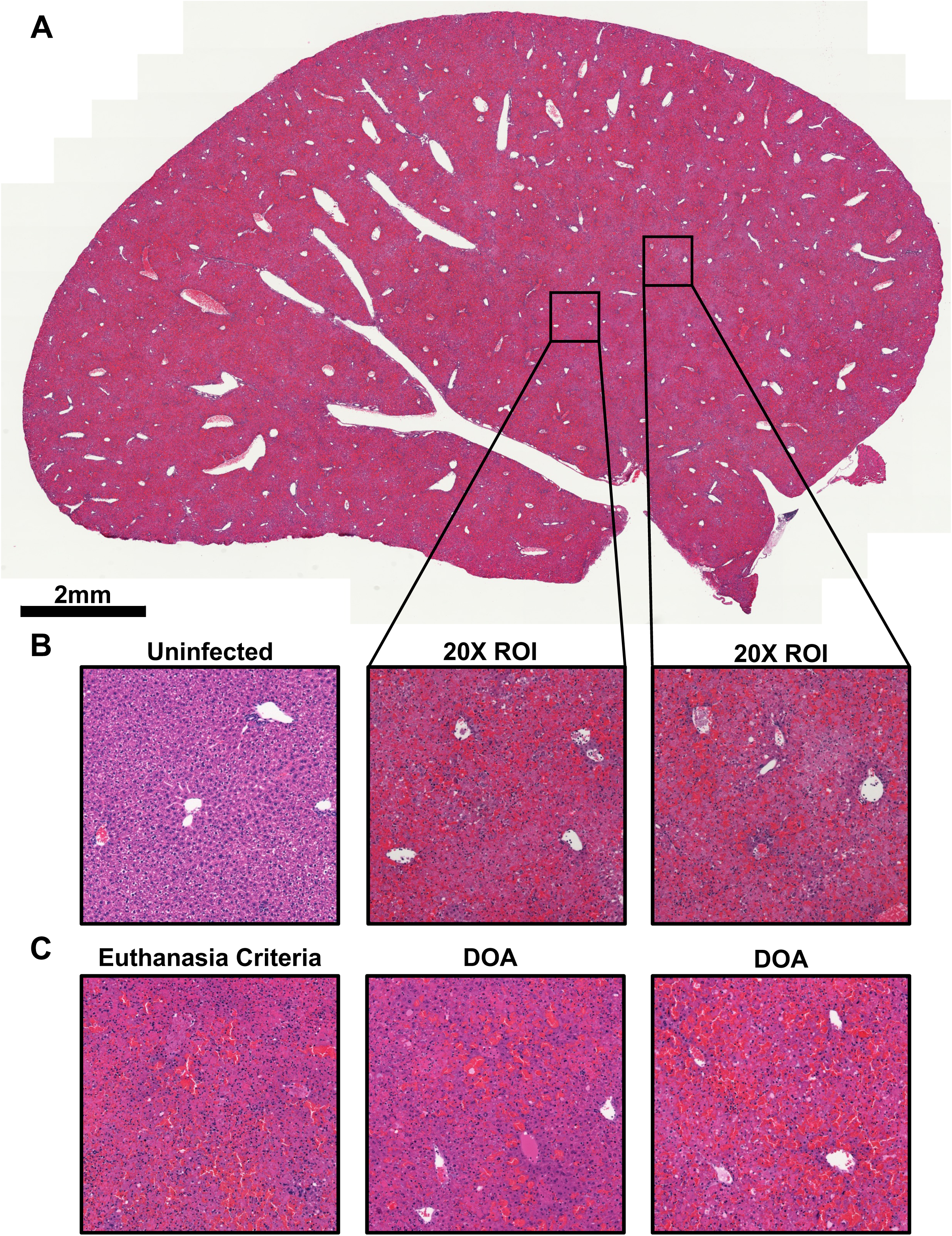
MAR1-treated mice have extensive hepatic damage at endpoint. (A) H&E staining of a representative FFPE liver left lateral lobe from a mouse treated with 500 µg MAR1 Ab 1 day prior to OROV infection with 10^3^ PFU and euthanized at a humane endpoint (total n=56). Composite of 20X images. (B) Representative H&E 20X ROIs (594×594 µm) from Figure 5A and an uninfected control. (C) Representative H&E 20X ROIs (594×594 µm) from 3 different mice at endpoint after infection with 10^6^ PFU OROV and 500 µg MAR1 treatment. Labels indicate whether the mouse was euthanized at a humane endpoint or found dead on arrival (DOA).

Given the severe disease and liver damage seen at endpoint (3-6 dpi across all doses), we treated another group of mice with MAR1 Ab and conducted a timed euthanasia study at 1, 2, and 3 dpi after footpad infection with 10^6^ PFU OROV. At each timepoint, blood chemistry, viral titer, and histopathology were assessed. In line with our previous results (Figure 4B), mice showed no clinical signs at 1-2 dpi, and 4/6 mice had succumbed to disease at 3 dpi. While there were no abnormal blood chemistry values at 1 dpi, by 2 dpi hepatocellular damage was observed with elevated ALT and normal ALP (Figure 6A, B; Supplementary Figure S9). At 3 dpi, 4 of 6 animals had succumbed to disease, therefore blood chemistry analysis was performed only on the remaining two mice. We found that these mice had ALT values in excess of 10,000 ALT U/L, high ALP levels, and a myriad of abnormal blood chemistry readings suggesting extensive liver damage and likely multiorgan failure (Figure 6A, B; Supplementary Figure S4). The blood chemistry values are in line with the massive hepatic necrosis seen at endpoint in our previous experiments (Figure 4). Abundant viral RNA was detectable as early as 1 dpi in the liver of euthanized mice and exceeded 10^8^ PFU/g equivalents by 2 dpi (Figure 6C). Infectious virus was detected in 3/6 livers at 1 dpi and increased to approximately 10^5^ PFU/g in all mice at 2 dpi (Supplementary Figure S10). At 3 dpi, while RNA titers further increased, infectious titers were lower (Supplementary Figure S10). Animals found dead on arrival or with extensive liver damage often have decreased infectious titers due to the high concentration of released liver enzymes and bile which inactivate infectious particles. It is likely that the extensive damage observed at 3 dpi is responsible for the fall in infectious titers at this timepoint while RNA titers marginally increased.

**Figure 6.**
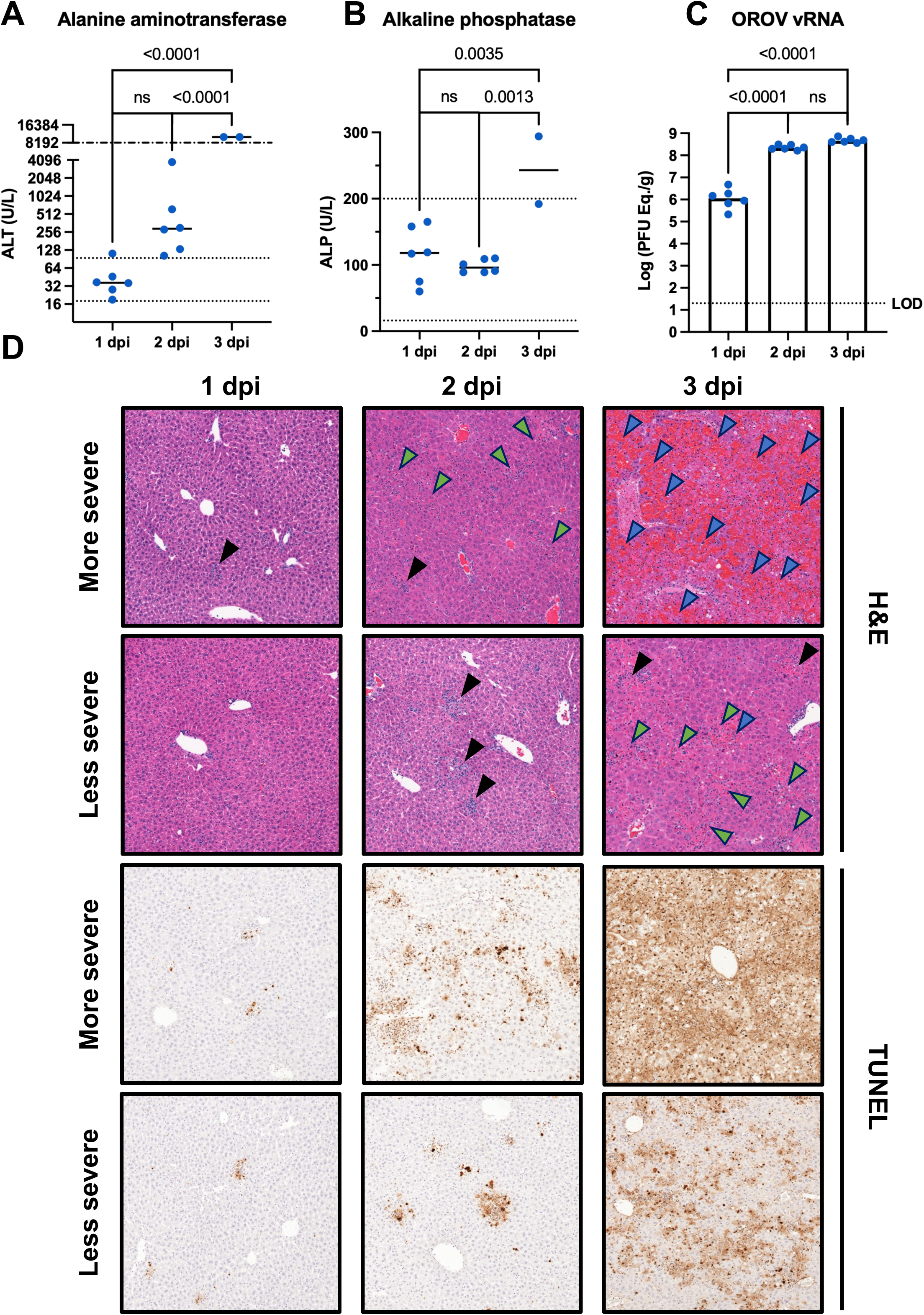
Liver injury in MAR1-treated mice is progressive and rapid onset. Mice were treated with 500 µg MAR1 Ab 1 day prior to infection with 10^6^ PFU OROV via footpad injection. A subset of animals was euthanized at 1 (n=6), 2 (n=6), and 3 (n=6) dpi. (A) Blood alanine aminotransferase (ALT) and (B) alkaline phosphatase (ALP) levels at indicated timepoints post infection. (C) Viral RNA in liver homogenate from tissues collected at the indicated timepoint quantified by RT-qPCR. Dotted lines indicate either normal (95 confidence interval) range or limit of detection (LOD). Dashed line with dots represents upper limit of the assay. (D) Representative H&E and TUNEL 20X ROIs (594×594 µm) from mice euthanized at 1, 2, and 3 dpi. Representative images have been partitioned into more severe or less severe based on pathologic interpretation. Black arrows mark necroinflammatory foci; green arrows mark bridging necrosis; blue arrows mark hemorrhage. Not all features are marked.

Histopathologic examination of whole liver left lateral lobes reveled that some animals displayed significant histopathologic damage (labeled “more severe”) while others had less prominent damage (“less severe;” Figure 6D; Supplementary Figure S11). This stratification aligns with the survival probabilities between 3-5 dpi where some animals are near death at 3 dpi while others may live to 5 dpi (Figure 4B). In more severe cases at 1 dpi, we found sparse necroinflammatory foci while less severe cases had no histopathologic signs of injury (Figure 6D). At 2 dpi, more severe cases had signs of sinusoidal congestion, coagulative necrosis, and lipid accumulation which is indicative of a lack of metabolic control due to extreme stress (Figure 6D). By contrast, the less severe cases at 2 dpi had necroinflammatory foci (Figure 6D). At 3 dpi, less severe cases mimic the more severe cases at 2 dpi; however, more severe cases present as a fulminant hepatitis with massive necrosis across the entire lobule as described above (Figure 4; Figure 6D). This variability is likely the reason why approximately 45% of mice at this dose would survive past 3 dpi. As in the acute hepatitis model, ALT levels correlate with TUNEL staining (Figure S12).

### Contemporary OROV outbreak isolate is less pathogenic in MAR1-treated mice

While the work with OROV BeAn19991 was ongoing, we received a more contemporary viral isolate (CDC 240023) from the ongoing outbreak; this isolate has been reported to be more pathogenetic in mice.^35^ OROV CDC 240023 is from the serum of a human patient who had recently traveled to Cuba.^18^ We evaluated the pathogenicity of OROV CDC 240023 in the lethal challenge model by treating mice with the MAR1 Ab one day before infection with 10^6^, 10^5^, or 10^4^ PFU. We found the contemporary CDC 240023 isolate caused markedly less lethality than the prototypical OROV strain (BeAn19991) across two independent experiments (Figure 7A). We also found that the time to death was delayed by approximately one day with a compressed clinical window (4-5 dpi for CDC 240023 versus 3-6 for OROV BeAn19991; Figure 7A). However, in the mice that succumbed to the contemporary OROV CDC 240023, we saw similar histopathology to the mice that succumbed to OROV BeAn19991, suggestive of fulminant hepatitis with massive hepatic necrosis (Figure 7B, 7C). While these results suggest that contemporary OROV strains may be less lethal in this particular mouse model, this does not negate the increased indices of severe outcomes seen in humans amid the ongoing outbreak. Rather, it suggests that current human isolates may be less adapted to infecting laboratory mice.

**Figure 7.**
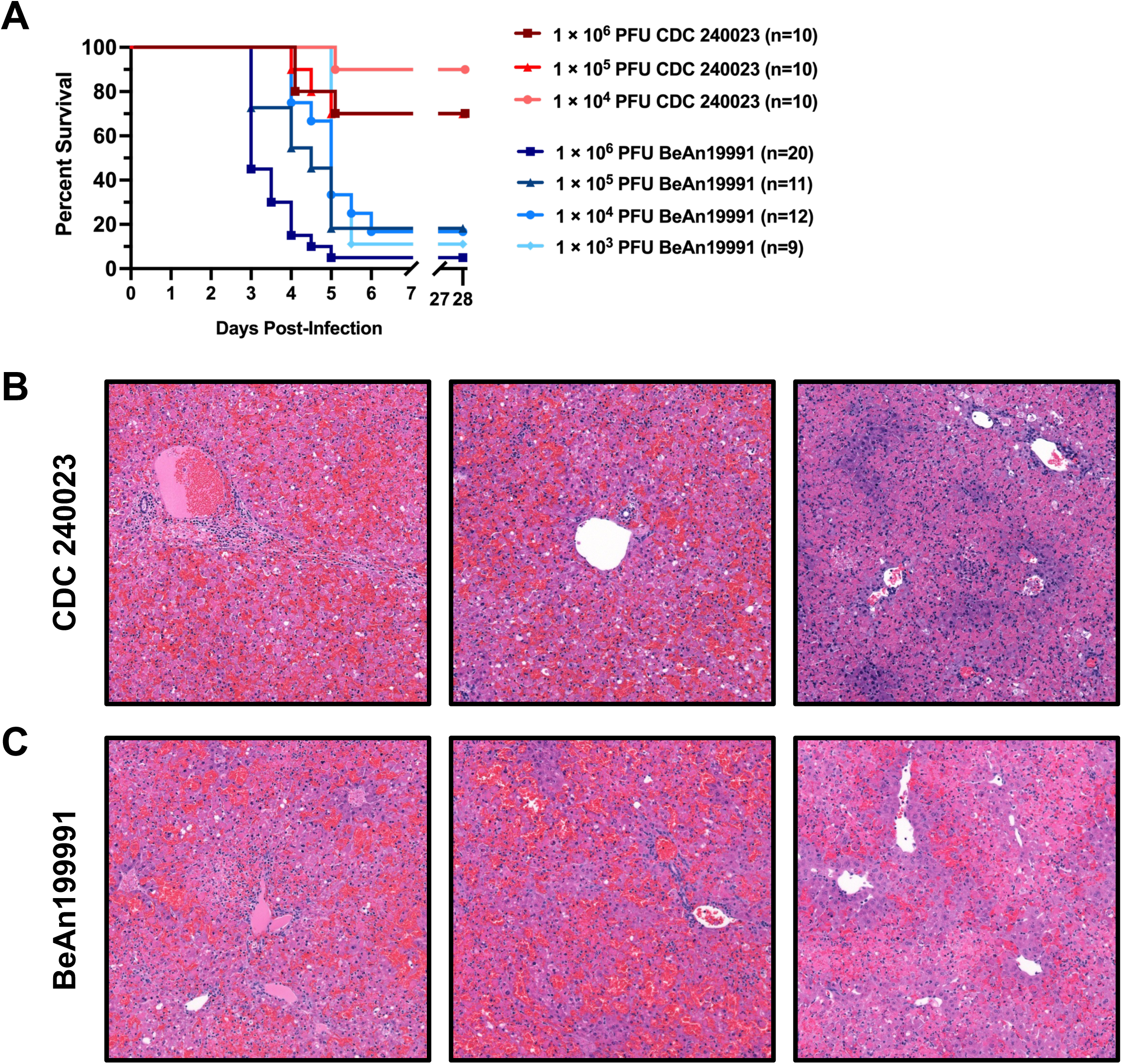
Contemporary OROV isolate is less pathogenic in laboratory mice than the prototypical strain. (A) Survival of mice treated with 500 µg MAR1 Ab 1 day prior to infection at the indicated OROV dose and strain via footpad injection. Representative H&E 20X ROIs from mice that succumbed to (B) the contemporary OROV CDC 240023 isolate (n=3) or (C) the recombinant prototypical OROV BeAn19991 (n=56).

## Discussion

Herein, we present two mouse models of OROV-induced hepatic disease. The first is a non-lethal model in immunocompetent mice which is characterized by transient, mild hepatocellular damage. The second is a lethal challenge model where mice are transiently immunocompromised using an anti-Ifnar1 mAb treatment. In both models, OROV infection is hepatotropic which aligns with recent reports of lethal OROV hepatic disease in humans. In combination, these two models may prove useful to understand why most OROV infections are controlled and mild, while a minority progress to severe coagulopathy and liver failure. Both models will be useful in pre-clinical therapeutic and vaccine testing.

Prior to the OROV outbreak which began in 2023, published mouse models of OROV infection relied on either young mice to increase susceptibility to neurologic disease^36,37^ or immunodeficient Type I IFN knockout (Ifnar^−/−^) mice to induce broad tissue susceptibility.^38,39^ Two recent reports did note OROV appeared hepatotropic in mice.^35,40^

The non-lethal acute hepatitis model characterized here opens avenues to explore OROV-induced hepatocellular damage and repair from both immunological and histopathological lenses. The identity, abundance, and function of infiltrating immune cells in the necrotic foci were not explored in this study but may shed light on the mechanisms of OROV-induced tissue damage, clearance, and hepatic repair. Additionally, further understanding of how the liver responds to focal infection may shed light on liver injury and repair more broadly. The lethal challenge model presented here is advantageous for preclinical vaccination studies as it allows vaccination in an immunocompetent state prior to lethal challenge. Additionally, this model can be applied to existing transgenic mouse lines without needing to breed them in immunocompromised backgrounds.

The similarity between ALT levels and histopathology of immunocompetent mice at 3 dpi and MAR1 treated mice at 2 dpi is striking. At these timepoints, we found mice have moderately elevated ALT levels ranging from the upper limits of the normal range to approximately 3500 ALT U/L. Mice also have similar histopathology with necrotic foci sharing a similar distribution and characteristics at these timepoints. While mice at these timepoints appear to be experiencing similar disease manifestations, in the immunocompetent mice the infection resolves and the liver repairs; whereas MAR1 treated mice progress to severe hepatic disease, necrosis, and death (Figure 8).

**Figure 8.**
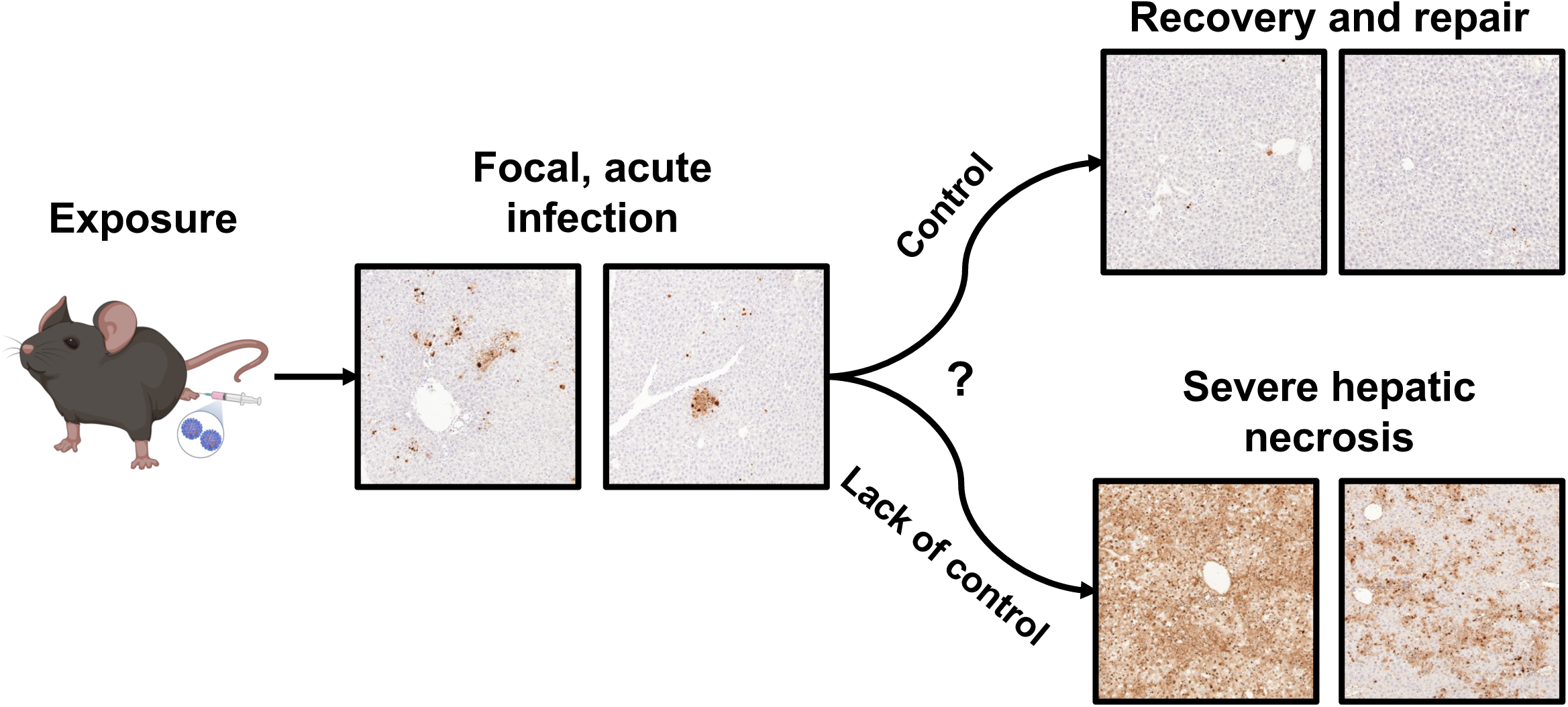
Model of liver pathogenesis caused by OROV. OROV infection in the footpad leads to focal infection of the liver. In immunocompetent mice, infection is controlled and repair follows (top). When the IFN response is suppressed, infection is uncontrolled and leads to massive necrosis and fulminant hepatitis (bottom).

While three lethal human cases of Oropouche fever provided detailed descriptions of liver histopathology, the third lethal human case also included representative microscopic images. The liver sections from this patient revealed severe hepatic damage, steatohepatitis, and necrotic foci which co-stained with OROV antigen, specifically in hepatocytes.^25^ This patient’s necrotic foci with infiltrating mononuclear cells looks remarkably similar to the inflammatory foci we found in the sub-lethal C57BL/6 model at 3 dpi and the MAR1-treated mice at 2 dpi. Considering the cause of death in this case was determined to be renal failure with respiratory distress (despite a low level of OROV antigen in these tissues), it makes sense that the level of necrosis in the liver is more similar to our immunocompetent mice or MAR1-treated mice prior to progression. Similarly, two recent human case reports from the current outbreak report liver enzyme elevation of a similar magnitude and pattern as we see in our models; that is, elevated ALT without elevation of ALP.^26,27^ The models presented here may provide insight into mechanisms of human OROV hepatic disease.

The single dose of MAR1 antibody that we used here was 5-fold below saturating level and yet it reliably induced OROV susceptibility. It is possible that lower doses of MAR1 may achieve the same outcome, or, conversely, higher doses with repeated maintenance doses may induce lethality at lower doses of OROV. Regardless, antagonizing the Type I interferon response with the MAR1 antibody leads to lethal disease, indicating that Type I interferons are a major OROV restriction factor, as previously reported.^38,41^ Additionally, antagonism of the Type I interferon system enables OROV to infect non-hepatic tissues at detectable levels, which suggests that OROV has mechanisms to evade the initial Type I interferon response specifically in the liver. By comparing these two models, we may be able to determine what causes different outcomes (recovery or progression to severe hepatis).

When comparing the prototype to a contemporary strain of OROV, the contemporary isolate OROV CDC 240023 was significantly less lethal in the MAR1 model than OROV BeAn19991, with an approximate 1 day lag in pathogenesis for the mice that succumbed. Similar results were also observed in a recently published study.^42^ In contrast, it was also recently reported that OROV CDC 240023 had 100% lethality in Ifnar^−/−^ mice with pathogenesis occurring 1 day earlier than BeAn19991.^35^

The discrepancy between these studies may be explained by the inocula used in each case. The BeAn19991 virus used here was generated by reverse genetics using the corrected sequence of the strain originally isolated in 1960.^43^ While recovered using reverse genetics, BeAn19991 likely had a rich passage history prior to sequences being deposited. It is likely this included suckling mouse brain passages, which may have mouse adapted BeAn19991 to some extent. In contrast, the CDC 240023 virus is the 4th passage of a recent clinical isolate from a human. It is not clonal and likely represents a mixed population. Further characterization of additional contemporary isolates, including with or without cloning and rescue, may be required to conclusively determine whether currently circulating strains are more virulent in mouse models. Additionally, lethality in a mouse model may not directly compare with virulence in humans.

In summary, the historically overlooked hepatotropic nature of OROV is now being appreciated due to the documented cases of lethal human disease with liver involvement as well as careful study of laboratory animals with and without innate immune blockade. Understanding more about how OROV causes liver infection in both a lethal and sublethal context will be useful for evaluation of vaccines and therapeutic treatments such as small molecule inhibitors or monoclonal antibodies.

## Methods

### Viruses

The prototypical strain of OROV (BeAn19991; GenBank accession numbers KP052850, KP052851, KP052852) was generated by reverse genetics as previously published and generously provided by Dr. Natasha Tilston-Lunel (Indiana University) and W. Paul Duprex (University of Pittsburgh).^44^ The stock used in this study represents the second passage in VeroE6 cells from the original rescue. The contemporary isolate of OROV (CDC 240023; GenBank accession numbers: PQ417950.1, PQ417949.1, PQ417948.1) was a generous gift from Brandy Russell (Arbovirus Reference Collection, Center for Disease Control and Prevention, Fort Collins, CO). The stock used in this study represents the fourth passage in VeroE6 cells (received at passage 2). OROV CDC 240023 was isolated from the serum a febrile human patient who contracted OROV in Cuba by amplification in VeroE6 cells.^18^ All viruses were propagated in VeroE6 cells in D2 media (DMEM supplemented with 2% FBS, 0.5 mg/mL Pen/Strep, 4.5 g/L glucose, 4 mM L-glutamine, 110 mg/L sodium pyruvate). Viruses were concentrated by ultracentrifugation at 107,000 x g for 3 hours using a Sorvall SureSpin 630 swinging bucket rotor in a Sorvall Discovery 90SE with a 25% sucrose cushion. Pelleted virus was resuspended overnight at 4 °C and then aggregates were removed by centrifugation at 3,000 x g for 10 minutes. Stocks were frozen in DMEM with a final concentration of 12% FBS, 0.5 mg/mL Pen/Strep, 4.5 g/L glucose, 4 mM L-glutamine, 110 mg/L sodium pyruvate, and 10 µM Tris Buffer (pH = 8.0). Viruses used to propagate concentrated stocks were sequence verified using the PCR-NGS method previously described.^45^ Stocks were titered by plaque assay.

### Animal Studies

All animal studies were conducted under Protocol 22102000, 24115814, or 25107233 adhering to the *Guide for the Care and Use of Laboratory Animals* and approved by the University of Pittsburgh Institutional Animal Care and Use Committee (IACUC). The University of Pittsburgh is fully accredited by the Assessment and Accreditation of Laboratory Animal Care (AAALAC). All mice used in these studies were from the C57BL/6J background. Experiments included both male and female mice to account for sex as a biologic variable. For antibody treatment, mice were intraperitoneally injected with 100 µL of sterile PBS containing 500 µg of either anti-Mouse IFNAR-1 [Clone MAR1-5A3], Purified in vivo GOLD™ Functional Grade (Leinco) or Mouse IgG1 GOLD, Clone hksp (Leinco). For infections, virus was diluted to the desired dose in D2 media. For footpad infections, 5–7-week-old mice were anesthetized with 2.5% vaporized isoflurane then inoculated intradermally (ID) in the left footpad (20 µL per mouse using 29 G needles and insulin safety syringes). For intranasal infections, 3–4-week-old mice were inoculated using 50 µL split between both nostrils. For survival studies, mice were scored daily then, if appropriate, euthanized at a human endpoint as defined by the IACUC-approved euthanasia criteria or recorded as dead on arrival (DOA). For timed-euthanasia experiments, mice were sedated with isoflurane prior to euthanasia. Blood was harvested by terminal cardiac puncture followed by harvesting of brain, liver, spleen, kidney, and lung tissue. Livers were dissected with the left lateral lobe being processed for histopathology, the right medial lobe being processed for immunofluorescent imaging, and the right lateral lobe and caudate process being homogenized for viral quantification. Tissues were homogenized in 0.5 mL D2 media using the Benchmark BeadBlaster 24 (6 m/s; 10 cycles; 30 sec. cycle/30 sec. pause).

### Virus Quantification

Viral stocks and viral load in tissues were quantified by plaque assay with a 3-day incubation. Viral load in tissue was also quantified by RT-qPCR quantified in PFU equivalents by generating a viral RNA standard curve from a stock of known titer. Homogenized tissue was diluted in D2 media for plaque assay or inactivated in TRIzol (Invitrogen) for RNA extraction and RT-qPCR. Exact procedures were previously described.^46^

### Blood Chemistry

Blood chemistry analysis was performed using the VetScan VS2 Chemistry Analyzer (Zoetis) with either the Comprehensive Diagnostic Profile or Mammalian Liver Profile disks. Whole blood was harvested by terminal cardiac puncture and placed into lithium heparin tubes; tubes were rocked until blood could be analyzed. In cases where ALT values exceeded the dynamic range of the analyzer (>2000 U/L), whole blood was diluted in 0.9% NaCl. In cases where insufficient blood was harvested or there were otherwise errors with the analyzer, those samples were excluded from presentation and analysis.

### Histopathology

Liver left lateral lobes were fixed in 10% neutral buffered formalin (NBF) then embedded in paraffin (FFPE) by the Cells, Tissues and Models Core (CTMC) previously Clinical Biospecimen Repository and Processing Core (CBRPC) at the Pittsburgh Liver Research Center (PLRC). The PLRC CTMC then performed hematoxylin and eosin (H&E) and TUNEL staining by standard methods. Microscopic histopathological interpretation was performed by Dr. Paul Monga (PLRC) blinded to the treatment groups. Brightfield images of stained slides were acquired using the Nikon PreciPoint Fritz Slide Scanner at the PLRC Advanced Cell and Tissue Imaging Core (ACTIC) within the Center for Biological Imaging (CBI; University of Pittsburgh) with a 20X Nikon CFI Plan Fluor objective. TUNEL staining was quantified using ImageJ by deconvoluting DAB pigmentation and thresholding TUNEL positive area over full tissue area. Thresholds were kept consistent across samples and experiments.

### Immunofluorescent Staining

Liver right medial lobes were fixed in NBF then cryoembedded in O.C.T. compound (Fisher). Cryosections (5 µm) were permeabilized with 0.1% Triton-X 100 made in PBS for 15 minutes. Tissues were washed with PBS and PBB (PBS supplemented with 0.5% BSA) then blocked with 5% Normal Goat Serum for 1 hour. A custom OROV N rabbit polyclonal antibody (GenScript) was diluted 1:200 in PBB and incubated with the tissue for 2 hours. Subsequently, Phalloidin 488 (Invitrogen) and a goat anti-Rabbit antibody conjugated with AlexaFluor 647 (Invitrogen) were applied at 1:400 and 1:1000 dilutions in PBB, respectively. Tissues were counterstained with Hoechst 33258 for 30 seconds and mounted using gelvatol provided by the PLRC ACTIC and CBI. Tissues were imaged using a Leica DMI8 inverted fluorescent microscope provided by the Center for Vaccine Research. All images were captured with the same settings and adjusted equally using ImageJ. A treated and untreated sample without primary antibody was used to adjust for background fluorescence.

### Statistics

For comparisons between blood chemistry values and viral titers, ordinary one-way ANOVA and Tukey’s multiple comparisons test were performed. Viral titers were log transformed prior to analyses and graphing. For comparison of ALT levels and TUNEL area, a simple linear correlation and Pearson’s correlation coefficient were computed with two-tailed P calculation. P values reported as exact or ns (not significant) if p > 0.05. Analyses were performed using GraphPad Prism (version 10.6.1).

### Data Availability

All data needed to evaluate the conclusions of this manuscript are presented in the manuscript and/or supplementary information. The data was stored in Microsoft Excel (version 16.106.2) and graphed using GraphPad Prism (version 10.6.1). Raw image files and full slide scans of histopathology are available upon request due to the large file size of each image.

## Acknowledgements

We thank Stacey Barrick for administrative support and undergraduate research assistant Gia Romano for additional support. We thank all members of the Hartman Lab for valuable discussion and feedback. We thank the Center for Biologic Imaging at the University of Pittsburgh for instrumentation support. We thank the Pittsburgh Liver Research Center at the University of Pittsburgh for tissue processing and pathology support. Figures 1A, 4A, and 8 were created in part using BioRender.

This work was funded in part by the following awards from NIH/NIAID: R01 AI178378 (ALH), R01 AI169850 (GKA/ALH), T32 AI049820 (CES & RER), and F32 AI191453 (RER). This work was also partially supported by NIH/NIAID award U19 AI181984 (PI SPJW) to ALH and UC7 AI180311 (PI WPD) to support the Operations of the University of Pittsburgh RBL within the Center for Virus Research. Support from the Pittsburgh Liver Research Center was funded in part by NIH/NIDDK P30 DK120531 (SPM). The funders had no role in study design, data collection and analysis, decision to publish, or preparation of the manuscript.

## Author Contributions

Author contributions were defined according to the NISO contributor roles taxonomy (CRediT) and are as follows: conceptualization (CES, RER, JJM, SPM, ALH), data curation (CES), formal analysis (CES), funding acquisition (WPD, GKA, ALH), investigation (CES, RER, JJM, BAS, AJD), methodology (CES, RER, JJM, SPM, ALH), project administration (CES, RER, ALH), resources (ALH), supervision (RER, ALH), validation (CES, RER, SPM, ALH), visualization (CES), writing – original draft (CES), writing – review and editing (CES, RER, SPM, ALH).

**Supplementary Figure S1. Susceptibility of immunocompetent mice to OROV infection after footpad or intranasal inoculation.** (A) Mice were infected with 10^6^ PFU OROV via footpad injection (n=6) and monitored 21 days for signs of disease. (B) Mean mouse weight (normalized as percent change from baseline; n=6) with standard deviation error bars. (C) Mice were inoculated with the shown dose of OROV in 50 µL intranasally and monitored 21 days for signs of disease. (D) Mean mouse weight (normalized as percent change from baseline; n=9 per group) with standard deviation error bars.

**Supplementary Figure S2. Blood chemistry at 1, 3, and 5 dpi following OROV infection.** Mice were infected with 10^6^ PFU OROV via footpad injection and a subset were euthanized at 1 (n=11), 3 (n=15), and 5 (n=11) dpi. Uninfected controls were also included (n=7). Each data point represents one mouse across three experiments; bars represent median value. Dotted lines indicate normal (95 confidence interval) range. Each analyte and relevant unit shown on individual graphs. Uninfected controls labeled “NI” (not infected).

**Supplementary Figure S3. Infectious titers and viral RNA at 1, 3, and 5 dpi following OROV infection.** Mice were infected with 10^6^ PFU OROV via footpad injection and a subset were euthanized at 1 (n=11), 3 (n=15), and 5 (n=11) dpi. (A) Infectious OROV titers by plaque assay or (B) viral RNA at indicated timepoints post infection and in indicated tissues. Each data point represents one mouse across three experiments; bars represent median value. For non-liver tissues, a subset (n=6 per timepoint) were quantified. Dotted lines indicate limit of detection (LOD) as defined by the tissue with the highest LOD.

**Supplementary Figure S4. TUNEL area correlates with blood ALT values at 3 dpi following OROV infection.** (A-C) TUNEL staining of representative FFPE liver left lateral lobes from mice euthanized at 3 days post infection (10^6^ PFU OROV inoculation dose; total n=15, 3 independent experiments) and concordant ALT values. Composite images and a representative 20X ROI (594×594 µm) are shown. (D) Correlation between normalized TUNEL area (TUNEL area over full tissue area) and ALT value per animal. Each data point represents one mouse across three experiments (n=15). Simple linear regression and Pearson’s correlation coefficient were computed.

**Supplementary Figure S5. Representative full lobe scans at 1, 3, and 5 dpi following OROV infection.** H&E and TUNEL staining of a representative FFPE liver left lateral lobe from a mouse euthanized staining at 1 (A; n=11), 3 (B; n=15), and 5 (C; n=11) dpi.

**Supplementary Figure S6. Representative full lobe scans of uninfected control mice.** H&E staining of representative FFPE liver left lateral lobes as full scans (A, C) or 20X ROIs (594×594 µm) from the same animal (B, D) with concordant ALT values.

**Supplementary Figure S7. Representative full lobe scans of uninfected control mice.** TUNEL staining of representative FFPE liver left lateral lobes as full scans (A, B).

**Supplementary Figure S8. Representative full lobe scans of MAR1 treated mice 28 dpi.** Mice were treated with 500 µg MAR1 Ab 1 day prior to OROV infection via intradermal footpad injection. H&E of a representative FFPE liver left lateral lobes and 20X ROIs (594×594 µm) are shown from mice which survived to 28 dpi.

**Supplementary Figure S9. Blood chemistry at 1, 2, and 3 dpi following OROV infection.** Mice were treated with 500 µg MAR1 Ab 1 day prior to infection at 10^6^ PFU OROV via intradermal footpad injection. A subset of animals was euthanized at 1 (n=6), 2 (n=6), and 3 (n=6) dpi. Each data point represents one mouse; bars represent median value. Dotted lines indicate normal (95 confidence interval) range. Each analyte and relevant unit shown on individual graphs.

**Supplementary Figure S10. Infectious titers and viral RNA at 1, 2, and 3 dpi following OROV infection with MAR1 treatment at -1 dpi.** Mice were treated with 500 µg MAR1 Ab 1 day prior to infection at 10^6^ PFU OROV via intradermal footpad injection. A subset of animals was euthanized at 1 (n=6), 2 (n=6), and 3 (n=6) dpi. (A) Infectious OROV titers by plaque assay or (B) viral RNA at indicated timepoints post infection and in indicated tissues. Each data point represents one mouse; bars represent median value. Dotted lines limit of detection (LOD).

**Supplementary Figure S11. Representative full lobe scans at 1, 2, and 3 dpi following OROV infection with MAR1 treatment at -1 dpi.** Mice were treated with 500 µg MAR1 Ab 1 day prior to infection at 10^6^ PFU OROV via intradermal footpad injection. A subset of animals was euthanized at 1 (A; n=6), 2 (B; n=6), and 3 (C; n=6) dpi. H&E (left) and TUNEL (right) staining of a representative FFPE liver left lateral lobe are shown.

**Supplementary Figure S12. TUNEL area and blood ALT values correlate at 2 dpi following OROV infection with MAR1 treatment at -1 dpi.** (A-C) TUNEL staining of a representative FFPE liver left lateral lobe from a mouse euthanized 2 days post infection (10^6^ PFU OROV inoculation dose; n=6) and concordant ALT values. Composite images and a representative 20X ROI are shown. (D) Correlation between normalized TUNEL area (TUNEL area over full tissue area) and ALT value per animal. Each data point represents one mouse (n=6). Simple linear regression and Pearson’s correlation coefficient were computed.

## References

1 Sakkas, H., Bozidis, P., Franks, A. & Papadopoulou, C. Oropouche Fever: A Review. Viruses 10, 175 (2018). 10.3390/v10040175

2 Mourão, M. P. G. et al. Oropouche Fever Outbreak, Manaus, Brazil, 2007–2008. Emerging Infectious Diseases 15, 2063–2064 (2009). 10.3201/eid1512.090917

3 Pinheiro, F. P. et al. Oropouche virus. I. A review of clinical, epidemiological, and ecological findings. Am J Trop Med Hyg 30, 149–160 (1981).

4 World Health Organization. Oropouche virus disease in the Region of the Americas. (World Health Organization, 2024).

5 Pan American Health Organization. Public Health Risk Assessment related to Oropouche Virus (OROV) in the Region of the Americas - 3 August 2024. (World Health Orangization, 2024).

6 Pan American Health Organization. Epidemiological Update Oropouche in the Americas Region - 11 February 2025. (World Health Organization, 2025).

7 Pan American Health Organization. Epidemiological Update Oropouche in the Americas Region - 13 August 2025. (World Health Organization, 2025).

8 Naveca, F. G. et al. Human outbreaks of a novel reassortant Oropouche virus in the Brazilian Amazon region. Nat Med 30, 3509–3521 (2024). 10.1038/s41591-024-03300-3

9 Sah, R. et al. Oropouche fever outbreak in Brazil: an emerging concern in Latin America. Lancet Microbe (2024). 10.1016/S2666-5247(24)00136-8

10 Martins-Filho, P. R., Carvalho, T. A. & Dos Santos, C. A. Oropouche fever: reports of vertical transmission and deaths in Brazil. Lancet Infect Dis 24, e662–e663 (2024). 10.1016/S1473-3099(24)00557-7

11 Castilletti, C. et al. Replication-Competent Oropouche Virus in Semen of Traveler Returning to Italy from Cuba, 2024. Emerg Infect Dis 30, 2684–2686 (2024). 10.3201/eid3012.241470

12 Control, E. C. f. D. P. a. Threat assessment brief: Oropouche virus disease cases imported to the European Union – 9 August 2024. (2024).

13 Morrison, A. et al. Oropouche Virus Disease Among U.S. Travelers - United States, 2024. MMWR Morb Mortal Wkly Rep 73, 769–773 (2024). 10.15585/mmwr.mm7335e1

14 Bello-Rodriguez, B. M., Vega-Jimenez, J., Canete, R. & Rodriguez-Morales, A. J. Emergence of Oropouche Virus Infection in Matanzas, Cuba, 2024. J Infect, 106470 (2025). 10.1016/j.jinf.2025.106470

15 Benitez, A. J. et al. Oropouche Fever, Cuba, May 2024. Emerg Infect Dis 30, 2155–2159 (2024). 10.3201/eid3010.240900

16 McGregor, B. L., Connelly, C. R. & Kenney, J. L. Infection, Dissemination, and Transmission Potential of North American Culex quinquefasciatus, Culex tarsalis, and Culicoides sonorensis for Oropouche Virus. Viruses 13 (2021). 10.3390/v13020226

17 Scroggs, S. L. P. et al. Enhanced infection and transmission of the 2022-2024 Oropouche virus strain in the North American biting midge Culicoides sonorensis. Sci Rep 15, 27368 (2025). 10.1038/s41598-025-11337-8

18 Payne, A. F. et al. Lack of Competence of US Mosquito Species for Circulating Oropouche Virus. Emerg Infect Dis 31, 619–621 (2025). 10.3201/eid3103.241886

19 Mancuso, E. et al. Assessing the vector competence of Italian Culex pipiens and Aedes albopictus mosquitoes for the re-emerging Oropouche virus. Parasit Vectors 18, 268 (2025). 10.1186/s13071-025-06912-x

20 de Armas Fernandez, J. R., et al. Report of an unusual association of Oropouche Fever with Guillain-Barre syndrome in Cuba, 2024. Eur J Clin Microbiol Infect Dis 43, 2233–2237 (2024). 10.1007/s10096-024-04941-5

21 Bandeira, A. C. et al. Fatal Oropouche Virus Infections in Nonendemic Region, Brazil, 2024. Emerg Infect Dis 30, 2370–2374 (2024). 10.3201/eid3011.241132

22 Ceccarelli, G. et al. Oropouche virus infection: Differential clinical outcomes and emerging global concerns of vertical transmission and fatal cases. Int J Infect Dis 150, 107295 (2025). 10.1016/j.ijid.2024.107295

23 Sah, R. et al. Oropouche fever fatalities and vertical transmission in South America: implications of a potential new mode of transmission. Lancet Reg Health Am 38, 100896 (2024). 10.1016/j.lana.2024.100896

24 Vernal, S., Martini, C. C. R. & da Fonseca, B. A. L. Oropouche Virus-Associated Aseptic Meningoencephalitis, Southeastern Brazil. Emerg Infect Dis 25, 380–382 (2019). 10.3201/eid2502.181189

25 Co, A. C. G. et al. Unravelling the pathogenesis of Oropouche virus. Lancet Infect Dis 25, e381–e382 (2025). 10.1016/S1473-3099(25)00341-X

26 Caetano Ferreira, K. G., Abdalla Santos, J. H., Ribeiro Pinheiro, G., Fernandes Abdalla, L. & Carneiro Dos Santos, L. Rhabdomyolysis and Hepatitis Associated with Oropouche Fever: Brazil. Am J Trop Med Hyg (2025). 10.4269/ajtmh.25-0107

27 Li, J. et al. Human liver-derived organoids recapitulate Oropouche virus infection and manifestation, enabling antiviral drug discovery. Cell Rep Med, 102646 (2026). 10.1016/j.xcrm.2026.102646

28 Otto, G. P. et al. Clinical Chemistry Reference Intervals for C57BL/6J, C57BL/6N, and C3HeB/FeJ Mice (Mus musculus). J Am Assoc Lab Anim Sci 55, 375–386 (2016).

29 Oh, B. et al. Difference in Intraspecies Transmissibility of Severe Fever with Thrombocytopenia Syndrome Virus Depending on Abrogating Type 1 Interferon Signaling in Mice. Viruses 16 (2024). 10.3390/v16030401

30 Xu, Z. S. et al. LDLR is an entry receptor for Crimean-Congo hemorrhagic fever virus. Cell Res 34, 140–150 (2024). 10.1038/s41422-023-00917-w

31 Park, S. C. et al. Pathogenicity of severe fever with thrombocytopenia syndrome virus in mice regulated in type I interferon signaling: Severe fever with thrombocytopenia and type I interferon. Lab Anim Res 36, 38 (2020). 10.1186/s42826-020-00070-0

32 Wilken, L., Stelz, S., Prajeeth, C. K. & Rimmelzwaan, G. F. Transient Blockade of Type I Interferon Signalling Promotes Replication of Dengue Virus Strain D2Y98P in Adult Wild-Type Mice. Viruses 15 (2023). 10.3390/v15040814

33 Sheehan, K. C. et al. Blocking monoclonal antibodies specific for mouse IFN-alpha/beta receptor subunit 1 (IFNAR-1) from mice immunized by in vivo hydrodynamic transfection. J Interferon Cytokine Res 26, 804–819 (2006). 10.1089/jir.2006.26.804

34 Dunn, G. P. et al. A critical function for type I interferons in cancer immunoediting. Nat Immunol 6, 722–729 (2005). 10.1038/ni1213

35 Gunter, K. B. et al. From prototype to outbreak: conserved pathogenesis of Oropouche virus in a novel murine pregnancy model highlights its public health implications. bioRxiv (2025). 10.1101/2025.08.02.668287

36 Santos, R. I. et al. Experimental infection of suckling mice by subcutaneous inoculation with Oropouche virus. Virus Res 170, 25–33 (2012). 10.1016/j.virusres.2012.07.006

37 Santos, R. I. et al. Spread of Oropouche virus into the central nervous system in mouse. Viruses 6, 3827–3836 (2014). 10.3390/v6103827

38 Proenca-Modena, J. L. et al. Oropouche virus infection and pathogenesis are restricted by MAVS, IRF-3, IRF-7, and type I interferon signaling pathways in nonmyeloid cells. J Virol 89, 4720–4737 (2015). 10.1128/JVI.00077-15

39 Gunter, K. et al. A reporter Oropouche virus expressing ZsGreen from the M segment enables pathogenesis studies in mice. J Virol 98, e0089324 (2024). 10.1128/jvi.00893-24

40 da Silva Menegatto, M. B., et al. Oropouche virus infection induces ROS production and oxidative stress in liver and spleen of mice. J Gen Virol 104 (2023). 10.1099/jgv.0.001857

41 Muraro, S. P. et al. Type I interferon controls vertical transmission and fetoplacental infection of Oropouche virus. iScience 29, 114647 (2026). 10.1016/j.isci.2026.114647

42 Yamada, Y. et al. Lineage-matched Oropouche virus mRNA-LNP vaccines confer complete, cross-protective immunity in mice. mBio, e0365525 (2026). 10.1128/mbio.03655-25

43 Acrani, G. O. et al. Establishment of a minigenome system for Oropouche virus reveals the S genome segment to be significantly longer than reported previously. J Gen Virol 96, 513–523 (2015). 10.1099/jgv.0.000005

44 Tilston-Lunel, N. L., Acrani, G. O., Randall, R. E. & Elliott, R. M. Generation of Recombinant Oropouche Viruses Lacking the Nonstructural Protein NSm or NSs. J Virol 90, 2616–2627 (2015). 10.1128/JVI.02849-15

45 Misu, M. et al. Rapid whole genome sequencing methods for RNA viruses. Front Microbiol 14, 1137086 (2023). 10.3389/fmicb.2023.1137086

46 Connors, K. A. et al. Neural cells are susceptible to historic and recently emerged Oropouche virus strains. PLoS Pathog 22, e1013933 (2026). 10.1371/journal.ppat.1013933

